# Task-specific, dimension-based attentional shaping of motion processing in area MT

**DOI:** 10.1101/058693

**Authors:** Bastian Schledde, F. Orlando Galashan, Magdalena Przybyla, Andreas K. Kreiter, Detlef Wegener

## Abstract

Non-spatial selective attention is based on the notion that specific features or objects in the visual environment are effectively prioritized in cortical visual processing. Feature-based attention (FBA) in particular, is a well-studied process that dynamically and selectively enhances neurons preferentially processing the attended feature attribute (e.g. leftward motion). In everyday life, however, behavior may require high sensitivity for an entire feature dimension (e.g. motion). Yet, evidence for feature dimension-specific attentional modulation on a cellular level is lacking. We here investigate neuronal activity in macaque motion-selective medio-temporal area (MT) in an experimental setting requiring the monkeys to detect either a motion change or a color change. We hypothesized that neural activity in MT is enhanced if the task requires perceptual sensitivity to motion. Despite identical visual stimulation, we found that mean firing rates were higher in the motion task, and response variability and latency were lower as compared to the color task. This task-specific response modulation in the processing of visual motion was independent from the relation between attended and stimulating motion direction. It emerged already in the absence of visual input, and consisted of a spatially global and tuning-independent shift of the MT baseline activity. The results provide single cell support for the hypothesis of a feature dimension-specific top-down signal emphasizing the processing of an entire feature class.

## Introduction

Neuronal processing of visual information strongly depends on the observer’s perceptual requirements (Chen et al. 2008; Huk and Heeger 2000; Jack et al. 2006). Among the several mechanisms involved in this selective, goal-directed processing, feature-based attention (FBA) facilitates the processing of a specific feature in the visual environment, independent of the spatial focus of attention, and at the cost of objects composed of different, irrelevant features (Bisley 2011; Maunsell and Treue 2006). Attending to a specific color hue, for example, elevates the neuronal response to an object at a remote, unattended location if its color matches the currently attended hue, but not if this object is composed of another, unattended hue (Saenz et al. 2002; Störmer and Alvarez 2014). Corresponding findings have been reported for attention directed to a specific motion direction, shape, or orientation (Corbetta et al. 1991; Liu et al. 2007; Saenz et al. 2002).

The same pattern is also observed at the level of single cells. Here, FBA is characterized by a firing rate (FR) change of neurons well-tuned to a specific, attended attribute within a feature dimension. For example, attending towards a specific motion direction globally increases the response of direction-selective MT neurons with a tuning preference for the attended direction (Treue and Martínez Trujillo 1999). The same neurons’ responses might be unmodulated (or even weakened) if attention is allocated on a motion direction that is significantly deviating from their preferred direction (Martínez Trujillo and Treue 2004). This feature-similarity gain (FSG), along with its spatially global effectiveness, has been suggested to underlie our ability to actually focus on the portion of the visual input that is behaviorally relevant, and, at the same time, disregard irrelevant, distracting information (Bisley 2011; Carrasco 2011; Maunsell and Treue 2006).

Under natural conditions, i.e. in a rapidly changing perceptual environment, successful behavior critically depends on the ability to quickly perceive abrupt changes of sensory input. In area MT, abrupt changes in motion induce strong, transient firing rate changes (Price and Born 2010; 2013), which in turn correlate with measures of perceptual performance (Galashan et al. 2013; Herrington and Assad 2009; Masse and Cook 2008; Traschütz et al. 2015). On the population level, these transients are carried substantially by neurons that are sub-optimally tuned to the stimulus feature preceding the feature-change (Traschütz et al. 2015) - hence, the population response relies significantly on neurons not addressed, or even suppressed, by FSG. We hypothesized, therefore, that under behavioral conditions requiring detection of changes in visual input, attentional facilitation is unlikely to be limited to the sub-class of neurons well-tuned to the currently attended feature attribute (which provide only minor information about the change), but will address all neurons processing information about the attended feature dimension, including those without a tuning preference for the currently attended feature attribute. Such tuning-independent, dimension-specific weighting of neural responses is supported by evidence from psychophysics (Found and Müller 1996; Müller et al. 1995), human EEG (Gledhill et al. 2015; Gramann et al. 2007; 2010; Pollmann et al. 2000; Töllner et al. 2008), and neuroimaging studies (Chawla et al. 1999; Pollmann et al. 2006; Weidner et al. 2002), but has not gained experimental support by single cell studies (Chen et al. 2012; Katzner et al. 2009), such that its underlying neuronal mechanisms remain unknown.

To address the notion of dimension-specific attention, we trained monkeys on two variants of a change detection task and performed extracellular recordings in area MT. In the first task, monkeys were required to detect speed changes - a feature MT neurons are highly sensitive to (Nover et al. 2005; Traschütz et al. 2015). In the second task, using identical visual stimulation, monkeys were required to detect color changes - a feature for which MT is only weakly sensitive (Croner and Albright 1999; Thiele et al. 1999). When comparing the neuronal representation to the same motion and motion change stimuli as a function of spatial attention and task requirements, we found that MT neurons were responding significantly different in the speed and in the color task, at the level of both evoked and spontaneous activity. Notably, this modulation was not only independent of the spatial focus of attention, but also independent of the relation between the attended and the preferred motion direction of a neuron, addressing neurons also when they were not tuned to the attended motion direction or speed. These results suggest a highly flexible, task-dependent shaping of motion processing.

## Material and Methods

### Electrophysiological recordings

All surgical and experimental procedures followed the Regulation for the Welfare of Experimental Animals issued by the Federal Government of Germany, and were approved by the local authorities. Extracellular recordings were obtained from two male adult rhesus monkeys *(Macaca mulatta)* using tungsten microelectrodes (0.8 - 5 MOhm, 125 µm shank diameter; Frederic Haer, Bowdoin, ME). Surgery was performed under Propofol/Fentanyl anesthesia and under strictly aseptic conditions, as previously reported in detail (Wegener et al. 2004). The recording chamber was placed over the middle temporal sulcus; coordinates for electrode penetrations were estimated from structural magnetic resonance imaging scans. During recordings, area MT was identified by the high proportion of direction selective neurons, the size/eccentricity ratio of RFs and the depth of the recording site (Desimone and Ungerleider 1986; Maunsell and Van Essen 1983; Mikami et al. 1986). The amplified electrode signal was sampled at a frequency of 25 kHz and band-pass filtered between 0.7 and 5 kHz, using either a custom-made hardware filter or an equiripple FIR filter in forward and reverse direction. Online detection of spikes was achieved by thresholding. At the beginning of each recording session, one or two electrodes were lowered through a guide tube penetrating the *dura mater* until the electrode’s tip reached the desired depth in area MT. Prior to cell recordings, the tissue was allowed to settle for about 30 min.

### Visual Stimulation

Visual stimuli were presented on a 22 inch CRT monitor (1280 × 1024 pixels, 100 Hz refresh rate), placed 83 cm from the animal. Stimuli were shown on a grey background (luminance: 10 cd/m^2^), and consisted of two high-contrast, drifting Gabors (spatial frequency: 2 cycles/deg), enveloped by a Gaussian with 0.75 deg at half height. Gabors had a mean luminance of 10 cd/m^2^ and drifted with 2.17 deg/sec. Speed and color changes were achieved by abruptly increasing the speed to 4.17 deg/sec or changing the color to an isoluminant pale yellow. Eye-movements were monitored with a custom-made eye-tracking device with a spatial resolution of 0.2 deg. Prior to cell recordings in the behavioral paradigms, monkeys performed a dimming task at fixation to determine basic response characteristics of each unit. RF size and location were mapped manually using a moving bar. If two electrodes were used simultaneously, we searched for units with largely overlapping RFs. Each units’ direction tuning was measured using Gabor gratings moving into one of 24 different directions.

### Behavioral task

Monkeys were trained to perform two variants of a feature-change detection task (Fig. 1A). They had to attend either inside or outside the RF or the recorded unit, and were required to detect either a speed or a color change. This 2*2 design allowed gathering data under the four experimental conditions illustrated by the inlet in Figure 2A. The monkey initiated a trial by maintaining fixation on a central fixation point (0.14° side length) and pressing a lever. The color of the fixation point indicated the task type (red: speed-change detection, yellow: color-change detection). 1050 ms after pressing the lever (or 350 ms for some recording sessions), a rectangular spatial cue indicating the location of the behaviorally relevant stimulus was displayed for 700 ms and outside the RF of the recorded neuron(s), followed by a 200 ms delay period. Subsequently, two static Gabor gratings appeared simultaneously. One grating was placed inside the RF and the other one at a mirror-inverted position in the opposite hemifield. 200 ms later, both gratings started to move intrinsically. If we recorded from two electrodes at the same time, the stimulus was placed in the joint RF of the units and motion direction of the RF stimulus was chosen to drive one of the recorded units with its preferred direction. Motion direction of the stimulus outside the RF was in opposite direction. If neurons had very different preferred motion directions, we chose a motion direction capable to drive both units efficiently. Thus, for a number of neurons we obtained data following stimulation with motion directions deviating to some degree from their preferred motion direction.

**Fig. 1.**
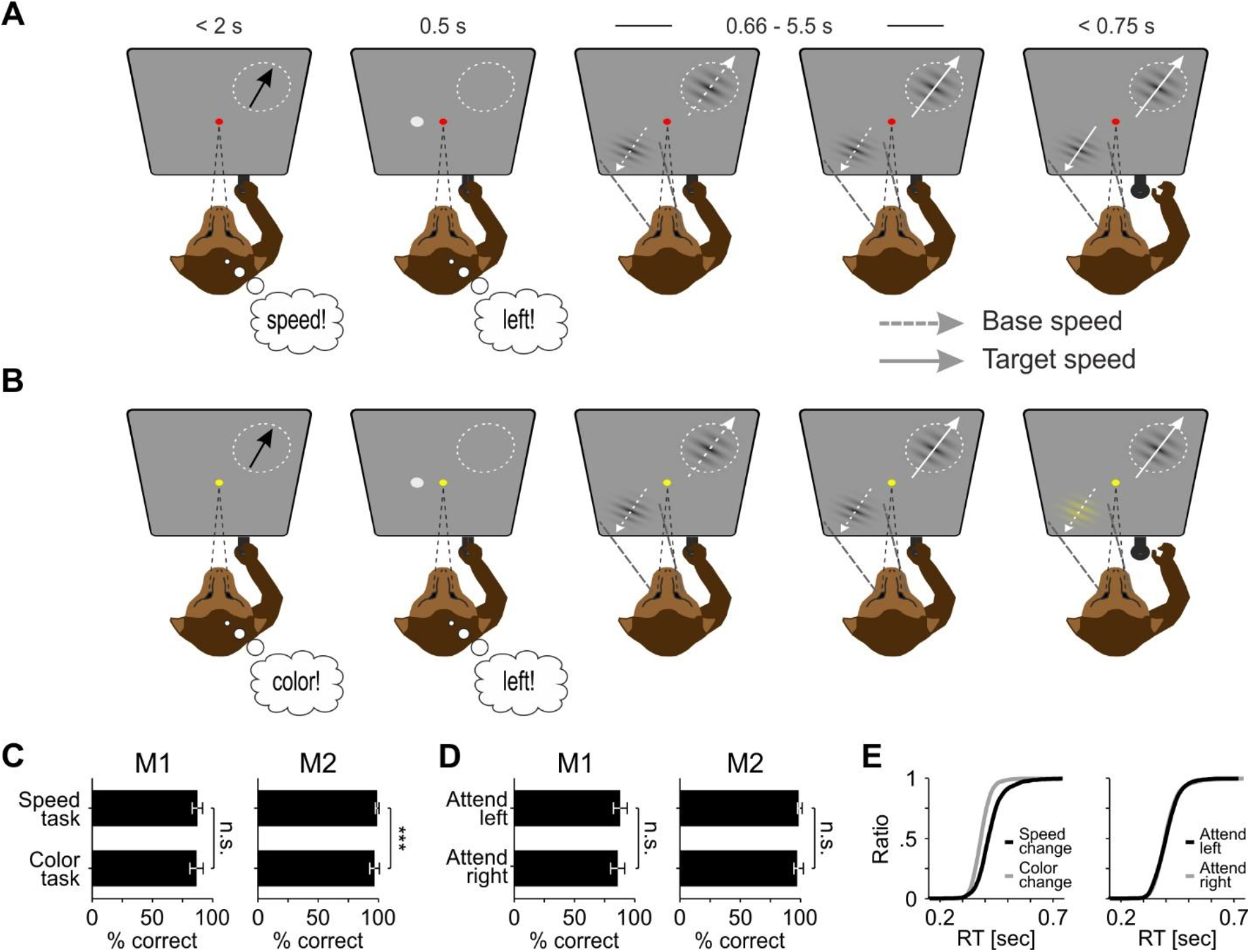
Behavioral paradigm and performance. *A* and *B*: Example trials of the speed-change task (*A*) and the color-change task (*B*). Task type was indicated by the color of the fixation point. Monkeys were required to detect a change of either the speed or the color at the cued location and to ignore any other change. Fixation had to be kept until 300 ms after releasing the lever (see Material and Methods for detailed information). Circle, RF of the recorded neuron; black arrow, preferred direction; White dashed arrow, motion direction at base speed; white straight arrow, motion direction at increased speed; thin dashed line, gaze direction; bold dashed line, direction of spatial attention. Note that visual stimulation is identical across tasks until the first change (here: RF stimulus speed-up, fourth panel from left in both tasks). *C* and *D*: Mean performance (correct responses/sum(correct responses, false alarms, misses)) of monkey M1 and M2 regarding task type (*C*) and spatial location of attention (*D*). *E*: Cumulative reaction time distributions of all trials, sorted by task type (left panel) and spatial allocation of attention (right panel).

**Fig. 2.**
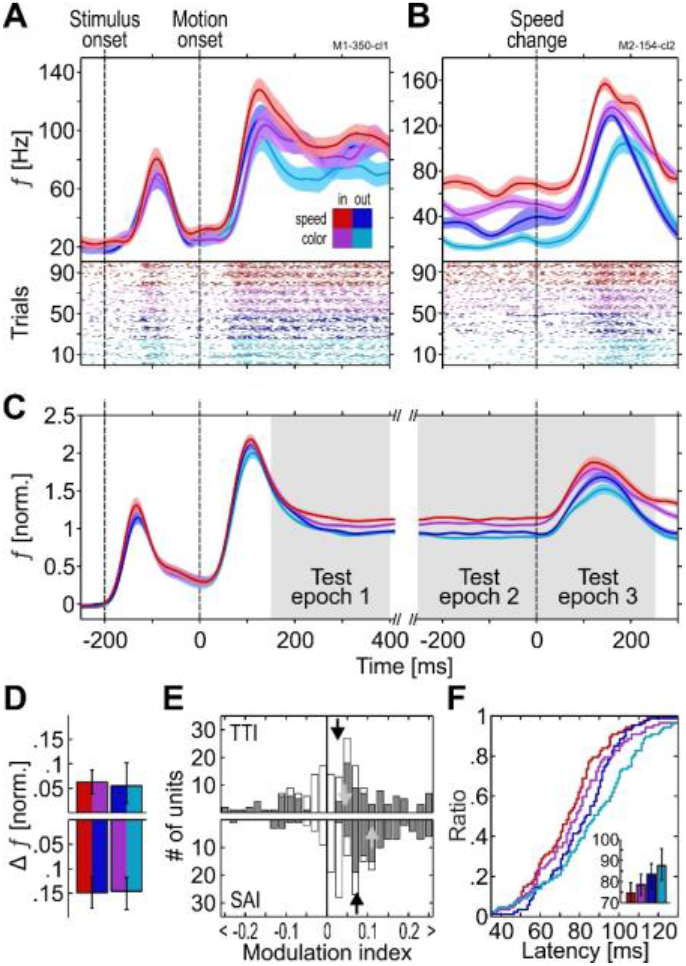
Task-specific modulation of firing rates. *A* and *B*: Spike density function (SDF) and corresponding scatter plot of two example units. Color scheme indicated by the inlet in (*A*) is used throughout the article. In, attend inside RF; out, attend outside RF; speed, detect speed change; color, detect color change. In the scatter plots at the bottom of the SDFs, each mark represents the time of a single spike. For illustration purposes, trials were sorted by condition. *C*: Population SDF (*N* = 187) during all task periods in each of the four conditions. Shaded areas: SEM. *D*: FR difference between conditions. Top, task effect; bottom, spatial attention effect. Left and right color in each bar indicate the attentional conditions that were subtracted from each other. *E*: Distribution of TTIs (top) and SAIs (bottom). Open bars, all units; gray bars, significantly modulated units. Black arrows, median AIs of all units; Gray arrows, median AIs of significantly modulated units. *F*: Cumulative distribution of response latencies during test epoch 3. Inlet: median latencies; error bars, 95% CI.

Following motion onset, monkeys had to attend the cued location and feature dimension and to signal detection of the target event (either a speed or a color change) by releasing the lever within 150 ms and 750 ms after the stimulus change. Target events occurred pseudo-randomly between 660 ms and 5500 ms after motion onset. Prior to these changes, up to three distractor events may have occurred, all of which had to be ignored. These distractor events consisted of a change in the currently relevant feature dimension at the uncued location, and/or a change in the irrelevant feature dimension at the cued and/or the uncued location. Their number and sequence varied pseudo-randomly. These trials increased the attentional demand for the monkeys and allowed us to verify that they were following the cue instructions. Throughout the trial and until 300 ms following the lever release, monkeys had to keep fixation within a circular eye window of 2 deg diameter, centered on the fixation point. To support selective attention to the relevant feature dimension (instead of more global attention to any change at the attended object), speed-changes and color changes were presented block-wise. The task order was alternated between recording sessions. We aimed to collect 25 successful trials of each experimental condition during which the speed change at the RF location was the first change event. This was behaviorally relevant only in the attend-in condition of the motion task, but irrelevant during the color task and the two attend-out conditions. Only these trials entered data analysis, because they allowed comparing the different attention conditions in response to identical visual stimulation. All other trials were disregarded.

### Data analysis

Data were analyzed using Matlab R2011b and later releases (The MathWorks, Natick, MA), using both in-built and custom-made functions. Spikes were detected and sorted semi-automatically as previously described in detail (Galashan et al. 2011). Spike thresholds were set by four times the median of the absolute values of the high-pass filtered signal (Quiroga et al. 2004), where applicable, otherwise by three times the signal’s SD. Occasionally, these thresholds had to be corrected manually. Preferred motion direction and significant direction tuning of the spike-sorted data were estimated offline using a response reliability approach (Grabska-Barwinska et al. 2012).

We collected data from 303 units, for which we obtained a sufficient visual response (> 1 SD above spontaneous activity and > 10 Hz) and at least 10 trials per experimental condition (mean: 25.1). 61 (20%) of these were excluded from data analysis due to slow electrode drifts over the course of the session. Electrode drifts were identified based on lab notes during recordings and careful visual inspection of the data, as well as by semi-automated procedures taking the mean FR over the course of the session into account. From the 243 remaining units, another 56 were not analyzed because they did not fulfill all of the following inclusion criteria: i) behavioral performance in both tasks above 75% (14%), ii) significant direction tuning (5%), iii) deviation between stimulating and preferred motion direction below 90 deg (3%). The final dataset consisted of 187 units gathered in 130 recording sessions. For 116 of these units, their preferred direction was within 15 deg of the stimulating motion direction (mean deviation: 7.1 deg), estimated by offline analysis of the spike-sorted data. For the remaining units, preferred and stimulating motion direction differed between 15.4 – 72.5 deg (mean: 30.9 deg, *N* = 71). All results derived from the entire database hold true if considering only units stimulated within 15 deg of their preferred motion direction.

Spike-density functions (SDF) were calculated within a 100 ms time window, shifted by 1 ms and smoothed by a Gaussian kernel (σ = 20 ms). Population responses were computed by subtracting each neuron’s mean baseline activity from its response, normalizing the response to the mean FR of all conditions during the period 400 ms prior to the speed change, and averaging over the normalized responses of all individual units. Modulation indices (MI) were defined as:

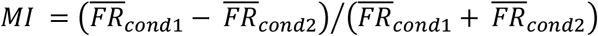

*Cond1* and *cond2* relate to each unit’s pool of trials from the speed-change and the color-change task, respectively, for investigating task-specific response differences (Task type index, TTI), and, analogously, to each unit’s pool of trials from the attend-in and attend-out conditions for investigating the effect of spatial attention (Spatial attention index, SAI).

Response latencies were calculated as the point in time at which the transient FR increase following the speed change reached 75% of its maximal peak (Galashan et al. 2013). Peak responses were required to occur between 20 and 300 ms after the speed change. Latencies were only calculated for units responding with a significant FR increase to the acceleration of the motion stimulus in each of the four experimental conditions (Wilcoxon signed rank, *α* < 0.05, *N* = 87). To investigate whether systematic differences in response latencies were independent from the magnitude of the transient speed-change response, we performed a rate-matching procedure by randomly searching for combinations of 10 trials per unit having the same transient peak rate in each of the two conditions to be compared (e.g. same peak rate in the speed and color task with attention allocated towards the RF). We found such trial combinations for 84 – 86 out of the 87 neurons with significant post-change transients in each experimental condition.

Trial-to-trial variability was measured by calculating the Fano factor (FF, mean normalized variance of spike counts) in successive, non-overlapping time windows during the 250 ms prior to the speed change. Window width varied between 5 ms and 125 ms, increased stepwise by 1 ms. For each width, the FF is reported as the mean of all windows. To investigate whether FF effects depended of FR differences between conditions, we performed another rate-matching procedure by searching for groups of 10 trials per experimental condition and unit for which the mean absolute FR did not differ by more than 1 Hz. This was done for successive, non-overlapping 50 ms windows during the 250 ms period starting before the speed change. For each of the windows we found such trial combinations for at least 174 of 187 units, and 182 neurons were considered for the statistical analysis of the pre-change epoch, during which FRs were relatively stationary.

Spike-count correlations (SCC) were computed for 38 pairs of simultaneously recorded units, using the same integration windows as for the FF. We only considered units recorded at different electrodes. For each pair, the raw spike counts were converted to z-scores, and the SCC was calculated as the Pearson correlation of z-scores across trials. Since this measure is sensitive to outliers, we excluded data segments which were above 3 SD of the mean of a given window and unit.

To investigate whether the response modulation by task or spatial attention condition depends on neuronal tuning, stimulus preferences of individual neurons were related to the actual stimulus properties. For direction preference, this was based on the deviation between stimulus direction and a unit’s preferred direction. For speed preference, we considered the transient FR change following the speed-up of the stimulus as an indicator for the match between stimulus and preferred speed. Because in area MT, most neurons are low-pass or follow a log-Gaussian shaped response profile (with preferred speeds in the range of base and target speeds as used in our experiments, or above) (Lagae et al. 1993; Nover et al. 2005), large transients unambiguously indicate neurons with a preferred speed clearly above the stimulating speed before the change (Traschütz et al. 2015). In turn, neurons with a small transient are likely to be driven by a speed that is more similar to their preferred speed, unless they have an extremely narrow or very broadband response profile. Thus, the size of the transient allows a reasonable approximation of how well the stimulus speed before the speed-change was matching the neuron’s preferred speed. The transient’s amplitude was calculated by dividing the mean FR during the 250 ms after the speed change by the mean FR during the 250 ms before the speed change, taken from the attend-in speed task. Direction preference-sorted neurons and speed preference-sorted neurons were distributed to groups of well-matching and less-matching neurons. For each subgroup (and for the individual neurons of the groups) demixed principal component analysis (dPCA) (Brendel et al. 2011; Kobak et al. 2014) was performed to quantify the amount of explained variance for the relevant response variables (time, spatial attention, task, and interaction of spatial attention and task). Demixed PCA was run using the original Matlab code as provided by the authors (https://github.com/machenslab/dPCA). In contrast to standard PCA, dPCA considers the data labels such that principal components depend on stimulus and task parameters, and allows demixing of data with arbitrary combinations of parameters. In our experiments, these parameters correspond to the stimulus-dependent neuronal response over time *T*, the two spatial attention conditions *SA*, and the two task conditions *TC.* The number of available data points *X* can be thought of as a *SA*-by-*TC* dimensional trajectory of length *T* in *N*-dimensional space, where *N* corresponds to the number of recorded units. By considering the labels of each data point, dPCA decomposes *X* into independent parts to estimate the variance attributable to one of the parameters *T*, *SA*, and *TC.* Thus, *X_T_* captures the variance due to stimulation over time, independent from spatial attention and task, while *X_SA_* captures the variance due to spatial attention that cannot be explained by *X_T_*. In turn, *X_TC_* describes the variance due to task conditions that cannot be explained by time and spatial attention condition, and *X_SA/TC_* describes the variance due to interaction between spatial attention and task condition that is not explained by the previous parameters. Since dPCA requires the same number of time bins *T* for all trials, the steady-state response between 200 ms after motion onset and 200 ms before the speed change was re-stretched to 260 time bins for each trial. For the shortest trials, this conformed to the original SDF resolution of 1ms/bin; for the longest trials each bin represents the SDF over a period of 4.4 ms. After re-stretching, each trial was described by a vector of 910 bins, lasting from motion onset to 250 ms after the speed change. To avoid overfitting, dPCA on each of the datasets was performed by adding a regularization term *λ*, as described in Kobak et al. (2014).

### Statistics

Statistical analysis was performed using non-parametric tests (balanced Friedman test, two-sided Wilcoxon signed rank test). Confidence intervals (CI) were obtained by bootstrapping the data with 5000 re-samples using the bias-corrected and accelerated percentile method (DiCicio and Efron 1996; Efron and Tibshirani 1993). For dPCA, periods of significant classification were computed by 100 iterations of stratified Monte Carlo leave-group-out cross-validation, as implemented in Kobak et al.(2014).

## Results

### Behavioral task and performance

Monkeys were trained on two variants of a feature-change detection task, and neuronal processing in area MT was investigated as a function of task requirements and spatial deployment of attention (Fig. 1, *A* and *B*). In both variants of the task, stimuli consisted of two grey Gabor patches with inherent motion. One Gabor was placed inside the RF of the recorded unit while the other one was inversely mirrored across the fixation point. The monkeys’ task was to detect either an instant 100% speed change or a sudden isoluminant color change to yellow. Each trial started with the appearance of a fixation point, indicating the task type by color (yellow: color task, red: speed task), followed by a spatial cue to signal target location (0.5 s). With a delay of 0.5 s, the two Gabors appeared simultaneously and started to move 0.2 s later. The Gabor within the RF moved approximately in the preferred direction of the recorded unit, and the other one moved in opposite direction. Color and speed changes occurred at pseudo-random points in time 0.66 s to 5.5 s after motion onset. They had to be ignored by the monkeys unless they were cued. The rationale of this 2*2-design was to implement two different task conditions that direct attention either towards or away from the motion domain, at a location inside or outside the neuron’s receptive field (RF). Speed- and color-change trials were presented block-wise and the order of blocks was balanced over recording sessions. Monkey 1 performed both tasks with 87%, with no statistical difference between tasks (Friedman test, *χ*^2^(1) = 0.88, *P* = 0.35, *N* = 53). Monkey 2 made extremely few errors and performed the speed and the color task with 99% and 97%, respectively, which nevertheless was statistically different [*χ*^2^(1) = 15.03, *P* < 10^−3^, *N* = 77) (Fig. 1*C*). For neither monkey there was a significant performance difference with respect to the attended location (both animals *χ*^2^(1) < 2.77, *P* > 0.096) (Fig. 1*D*). Regarding reaction times, monkeys detected color changes significantly quicker than speed changes (*χ*^2^(1) = 121.94, *P* < 10^−27^, *N* = 130), with color change detection leading by 31 ms on average (Fig. 1*E*). This is to be expected, due to the faster sensory processing of color, and in line with the processing time differences previously described (Arnold et al. 2001; Linares and López-Moliner 2006; Moutoussis and Zeki 1997). Attending towards or away from the RF yielded no significant difference between RT distributions (*χ*^2^(1) = 0.05, *P* = 0.82). Overall alertness was tested by calculating the time interval between fixation point-onset and trial initiation by lever press, assuming that increased or reduced alertness would lead to accelerated or delayed trial initiation, respectively. M1 initiated trials equally fast in both tasks (488 vs. 475 ms, *Z* = 1.17, *P* = 0.117), while M2 was slightly slower in the speed task (631 vs. 610 ms, *Z* = 2.86, *P* = 0.004). No changes in trial initiation time were observed during the course of the recording sessions (linear regression fits for both monkeys and tasks, all *F* < 3.61; all *P* > 0.06). Taken together, the behavioral data show no, or only minor, non-systematic differences regarding task conditions and spatial attention locations, indicating that both types of tasks and spatial attention conditions were about equally demanding for the monkeys.

### Task-specific firing rate modulation

We investigated task-related modulations of MT responses by comparing neuronal activity depending on whether motion was relevant or irrelevant for the task. Visual stimulation was identical across tasks until the first feature change (Fig. 1, *A* and *B*), such that for each of the two spatial attention conditions neuronal activity is expected to be the same in the speed and the color task, unless dependent on task requirements. Specifically, for attention directed to the RF, the motion direction of the attended Gabor matched, or was close to, the preferred direction of the recorded neuron. If FBA spreads to task-irrelevant target features (Katzner et al. 2009), the recorded neuron should receive about the same attentional gain in both tasks. In turn, for attention directed to the stimulus outside the RF, the motion direction of the attended Gabor is about 180 deg away from the stimulating, preferred motion direction of the recorded neuron. Under this condition, the neuron should receive no motion direction-related gain in any of the tasks (Martínez Trujillo and Treue 2004). Yet, for many of the recorded neurons we found a significant response difference between tasks, both with attention directed towards or away from the RF. Figure 2 provides two examples. For the single unit shown in Fig. 2*A* (with responses aligned to motion onset), recorded from monkey M1, FRs were considerably higher when the monkey performed the motion task as compared to the color task, independent of the spatial focus of attention. Likewise, the multi-unit shown in Figure 2*B* (with responses aligned to the speed change), recorded from monkey M2, was more active in the speed task throughout the period before the speed change as well as during the transient response following the speed-up of the RF stimulus. In all trials contributing to these SDFs, the speed change was the first change event, i.e. visual stimulation was identical across the four experimental conditions.

These units are representative of the population of MT neurons (187 units, M1: 24 single units, 46 multi-units, M2: 57 single units, 60 multi-units), calculated as the mean of the normalized, baseline-corrected responses of individual units. In both spatial conditions, FRs were consistently higher in the speed task than in the color task (Fig. 2*C*). This task-specific response difference was of smaller magnitude than the effect of spatial attention, but evident throughout the trial, affecting the steady-state response during stimulation with base speed as well as the transient response representing the speed change. Neurons responded strongest during the attend-in condition of the speed task and weakest during the attend-out condition of the color task. Statistical testing was carried out by a non-parametric ANOVA applied to the collapsed data of three 250 ms test-epochs (grey-shaded areas in Fig. 2*C*). Both the factor task and the factor spatial attention had a highly significant influence on the FR of the MT population (Friedman tests, task: *χ^2^*(1) = 11.85, *P* < 10^−3^; spatial attention: *χ^2^*(1) = 86.59, *P* < 10^−19^). Individual testing of the single epochs confirmed this finding (Friedman tests, all *χ^2^*(1) > 5.34, all *P* < 0.0209). To test whether the task effect was spatially global or, alternatively, restricted to the spatial focus of attention, we compared the firing rates between the two tasks and the two spatial attention conditions separately (Fig. 2*D*). Besides a spatial attention effect in both tasks (Wilcoxon signed rank test: both *Z* > 8.81, *P* < 10^−17^) and a task effect for the attend-in condition *(Z* = 3.78, *P* < 10^−3^), we found significantly higher FRs in the speed task for attention directed outside the RF (Z = 3.12, *P* = 0.0018), also if restricted to the mere population of single units (Z = 3.23, *P* = 0.0012, *N* = 81). The results hold also true for the individual animals (M1: all *Z* > 2.15, all *P* < 0.0313, *N* = 70; M2: all *Z* > 2.31, all *P <* 0.0206, *N =* 117). Thus, neuronal responses where consistently higher in the speed task, even when the attended motion direction was opposite to the preferred motion direction of the recorded neurons.

We next investigated how many of the individual units carried the task-specific response difference seen in the population. We found that 55% of all units had significantly different FRs between tasks (Friedman tests, *P* < 0.05), and 71% of these were more active in the speed task. To further test the sign and strength of this modulation, we converted the firing rate differences into a modulation index for task type-dependent response differences (TTI), ranging from −1 to 1. The TTI is positive for neurons having higher activity during the speed task (see Material and Methods). Collapsed over the three 250 ms time epochs, the median TTI of the entire population was 0.031 (significantly modulated units only: 0.059), and the distribution of index values was significantly greater than zero (Wilcoxon signed rank test, *Z* = −3.65, *P* < 10^−3^) (Fig. 2*E*, upper panel). The median TTI of the individual epochs 1 - 3 was 0.011, 0.037, and 0.03, respectively, and all distributions were significantly different from zero (Z < −2.3, *P* < 0.0217). We also calculated a spatial attention index (SAI) for comparison. 57% of all neurons were significantly modulated by spatial attention, and 92% of these were more active during the attend-in conditions. The collapsed median SAI of all neurons was 0.07 (significantly modulated units only: 0.106) (Fig. 2*E*, lower panel), and the median SAI of the individual epochs was 0.049, 0.075, and 0.076. All distributions were significantly different from zero (*Z* < −6.74, *P* < 10^−10^).

### Task-dependence of neuronal latencies

As seen from Figure 2, *B* and *C*, firing rate increments following the speed change of the RF stimulus not only differed in amplitude, but also seemed to vary in latency. Recently, we and others have shown that these latencies are significantly modulated by spatial attention (Galashan et al. 2013; Khayat and Martinez-Trujillo 2015; Sundberg et al. 2012) and closely correlate with reaction times (Galashan et al. 2013; Traschütz et al. 2015). Hence, because psychophysically reaction times for speed changes are faster in a speed task than in a color task (Wegener et al. 2008; 2014), we hypothesized that MT responses to speed changes will depend on task conditions as well, and be more rapid if speed is behaviorally relevant. We therefore analyzed the latency of the transient FR increase in response to the speed change (test epoch 3, cf. Fig. 2 *C*) of all neurons for which their speed tuning promoted an identifiable response peak in each of the four conditions (*N* = 87). Plotting of the cumulative RT distributions confirmed our previous result on the influence of spatial attention (Galashan et al. 2013), and also revealed a leftward-shift of the curves in the speed task both with attention directed to the RF and away from it (Fig 2*F*). In the attend-in conditions of the speed and the color task, the median latencies were 75 ms and 79 ms, respectively, and in the corresponding attend-out conditions, they were 84 ms and 88 ms, respectively (Fig. 2*F*, inlet). A Friedman test on the factors task and spatial attention revealed a significant influence of each factor (task effect: *χ*^2^ = 5.08, *P* = 0.0242; spatial attention: *χ*^2^ = 14.3, *P* < 10^−3^). *Post-hoc* Wilcoxon signed rank tests confirmed the task effect both with attention directed towards the RF and away from it, and also confirmed the influence of spatial attention on MT neuronal response latencies for both tasks (all *Z* > 2.14, all *P* < 0.0326). To further investigate whether this latency reduction is a native effect of attention, or, alternatively, a consequence of higher firing rates during the post-change response, we rate-matched transients of the four experimental conditions (see Material and Methods). Yet, latencies in the four comparisons (task effect inside and outside RF, spatial attention effect in speed and color task) were still significantly different (Wilcoxon signed rank tests, all *Z* > 1.97, all *P* < 0.04, *N* = [84 85 86 85]). This result indicates that, like spatial attention, task-dependent attentional modulation forces a leftward shift of the latency distribution within the speed task, independent of differences in the magnitude of the speed change response.

### Task requirements and neuronal variability

Because the signal-to-noise ratio of a population of neurons depends on both signal strength and variability, we next investigated neuronal response fluctuations. We calculated the Fano factor (FF) of each unit, and the spike-count correlation (SCC) of 38 simultaneously recorded pairs of units. Because independent and correlated response variability fluctuates on a range of time scales, both FF and SCC depend on the width of the integration window over which spikes are counted (Smith and Kohn 2008). To obtain an unbiased estimate, we used multiple window sizes (5 ms to 125 ms bin width, increased stepwise by 1 ms).

We applied this analysis to the period 250 ms prior to the speed change (while FRs were relatively stationary). Both the FF and SCC varied significantly as a function of behavioral condition (Fig. 3, *A* and *C*). For example, for a window size of 50 ms we found significantly different FFs between tasks both for attention inside and outside the RF (Wilcoxon signed rank test, *Z* > 2.29, *P* < 0.0222, *N* = 187). Likewise, the SCC of both spatial conditions was significantly smaller in the speed task (*Z* > 2.94, *P* < 0.0033, *N* = 38). Testing other window sizes demonstrated that the statistical outcome of the FF was consistent among most widths below 100 ms, indicating a reduced trial-to-trial variability in the speed task (Fig. 3*B*). For the SCC, statistical differences were found for the majority of integration windows, independent of the spatial focus of attention (Fig. 3*D*). For comparison, we also calculated the effect of spatial attention and found a corresponding and very robust influence, in line with previous results (Cohen and Maunsell 2009; Cohen and Newsome 2008; Galashan et al. 2013). The effect on the FF was highly significant (*Z* > 5.1, *P* < 10^−4^) for all window sizes, and SCC results were consistent among most windows (Fig. 3, *B* and *D*). Spatial attention had a stronger influence on the FF than the task type, but their effect on the SCC was about equally strong. We also tested whether the FF modulation was mainly due to differences in FR, or alternatively, independent from these. We applied a mean-matching procedure to five non-overlapping 50 ms time windows and estimated the FF for an integration window of 50 ms. FFs were smaller during the speed task (Friedman test, *χ*^2^ = 4.77, *P* = 0.029, *N* = 186) and for attention directed to the RF (*χ*^2^ = 33.21, *P* < 10^−8^). *Post-hoc* tests revealed a significant task effect inside the RF (Wilcoxon signed rank, *Z* = 2.37, *P* = 0.0177), but not so for attention directed away (*Z* = 1.08, *P* = 0.2821). Spatial attention had a significant effect in both tasks (*Z* > 4.22, *P* < 10^−4^). Thus, reductions in trial-to-trial variability occur independent of changes in FR for both task-specific and spatially attended information, yet presumably restricted to the attended location.

**Fig. 3.**
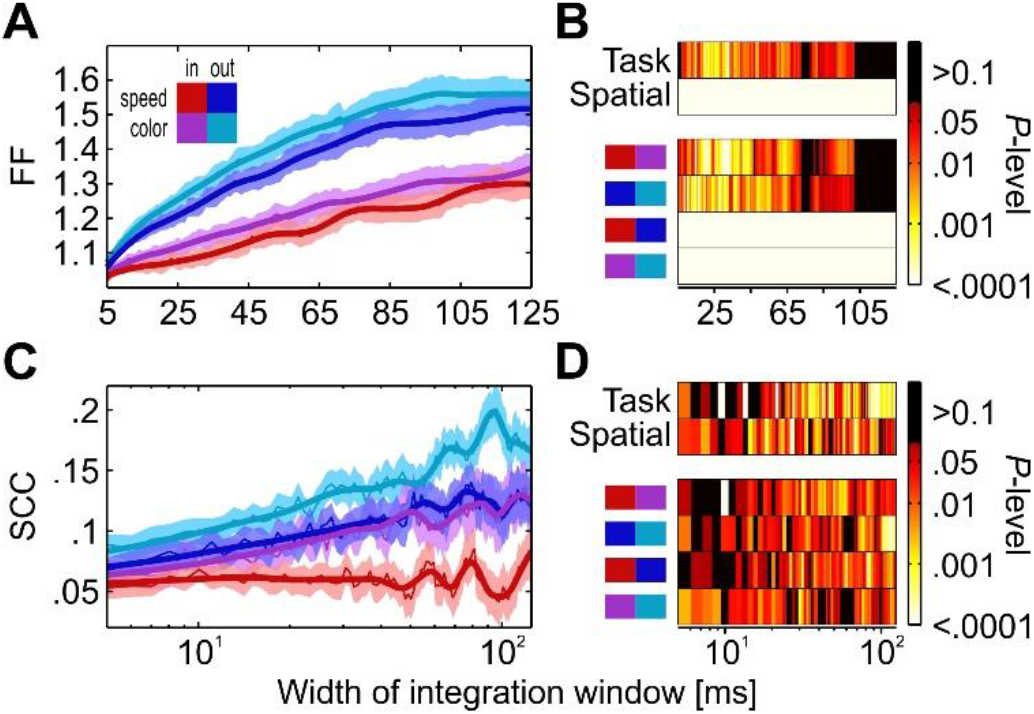
Variability of responses depending on attentional condition. *A* and *B*: Fano Factor of spike counts for integration windows between 5 ms and 125 ms width (*A*), and color-coded P-values for Friedman tests (top rows) investigating the factors task and spatial attention, and Wilcoxon tests (bottom rows) comparing single conditions (*B*). Color coding ranges between P-values of 10^−4^ and 10^−1^. Larger and smaller values are colored black and white, respectively. *C*: Spike-count correlation of 38 pairs of simultaneously recorded neurons for different integration windows. *D*: Statistical outcome for different integration window widths. Conventions as in *B*.

### Relation between neuronal tuning and magnitude of response modulation

The various analyses described earlier provide evidence for a robust influence of the specific change detection requirements on various parameters of the neuronal population response in MT. To further test this, we next asked whether the strength of task-dependent modulation relates to the neurons’ specific tuning properties, or alternatively, is independent from these. Since stimulus direction sometimes deviated from the neurons’ preferred direction (mostly due to paired recordings) and stimulus speed was not matched to the neurons’ speed tuning properties, we investigated the size of the task effect as a function of these deviations. To this end, we sorted all neurons according to i), the difference between their preferred motion direction and the actual stimulus direction (Fig. 4*A*), and ii), the size of the transient following the speed change (as an indicator of the match between stimulus base speed and a neuron’s preferred speed, see Material and Methods) (Fig. 4*C*). Based on this, we distributed all units into two groups of well-matching (*N* = 94) and less-matching (*N* = 93) preference, either by direction or by speed (Fig 4, *B* and *D*). We then performed demixed PCA analysis (Brendel et al. 2011; Kobak et al. 2014) on these groups to investigate how much variance of the response can be explained in terms of task parameters (see Material and Methods). Demixed PCA (dPCA) seeks to capture the maximum amount of variance by a minimum number of parameters, and allows to reconstruct the time course of spike trains as a function of these parameters (Brendel et al. 2011). Based on 20 components, dPCA explained between 81% and 90% of the total variance in the four groups, which was slightly below standard PCA (95% - 97%). For all groups, most of the variance (75 – 84%) was explained by the stimulus’ time course (from motion onset to 250 ms following the speed change), independent of condition (Fig. 4, *E* - *H*). Spatial attention explained between 7 and 10% of the variance, and task conditions explained another 7 to 11%. The remaining part of explained variance was due to the interaction between spatial attention and task (2 – 4%). Reconstructed neuronal responses (Fig. 4, *I* – *X*) were different regarding the condition-independent component of well-matching units when data were sorted according to the putative speed preference (due to a missing deflection of their response following the speed change, Fig. 4*L*), but where otherwise very similar between groups. Independent of the sorting criteria, spatial attention was significantly classified after the firing rate reached a steady state level, about 200 ms after motion onset, and during the remaining response period (Fig 4, *M*–*P*). Explained variance by task type allowed for significant classification of the task during the entire trial period, starting already during the initial motion onset transient and lasting until the end of the response (Fig. 4*Q*–*T*). This difference in time course for classifying the spatial attention and task conditions may solely reflect the difference in cueing (task condition lasted for a block, spatial attention condition was cued at the beginning of each trial). Rather, what the result of this analysis suggests is that the amount of explained variance by task type is independent of whether the neurons are sorted according to their speed or direction preference, or whether they are well-tuned or less-tuned to that feature. Thus, the group-wise analysis provides evidence for an independence of the task effect on the tuning properties of the neurons, at least within the range of deviations between preferred and stimulus features we covered. In terms of explained variance, task and spatial attention had about equally strong effects on neuronal responses in each of the groups, arguing for a dimension-based modulation of response properties due to the specific demands of the task.

**Fig. 4.**
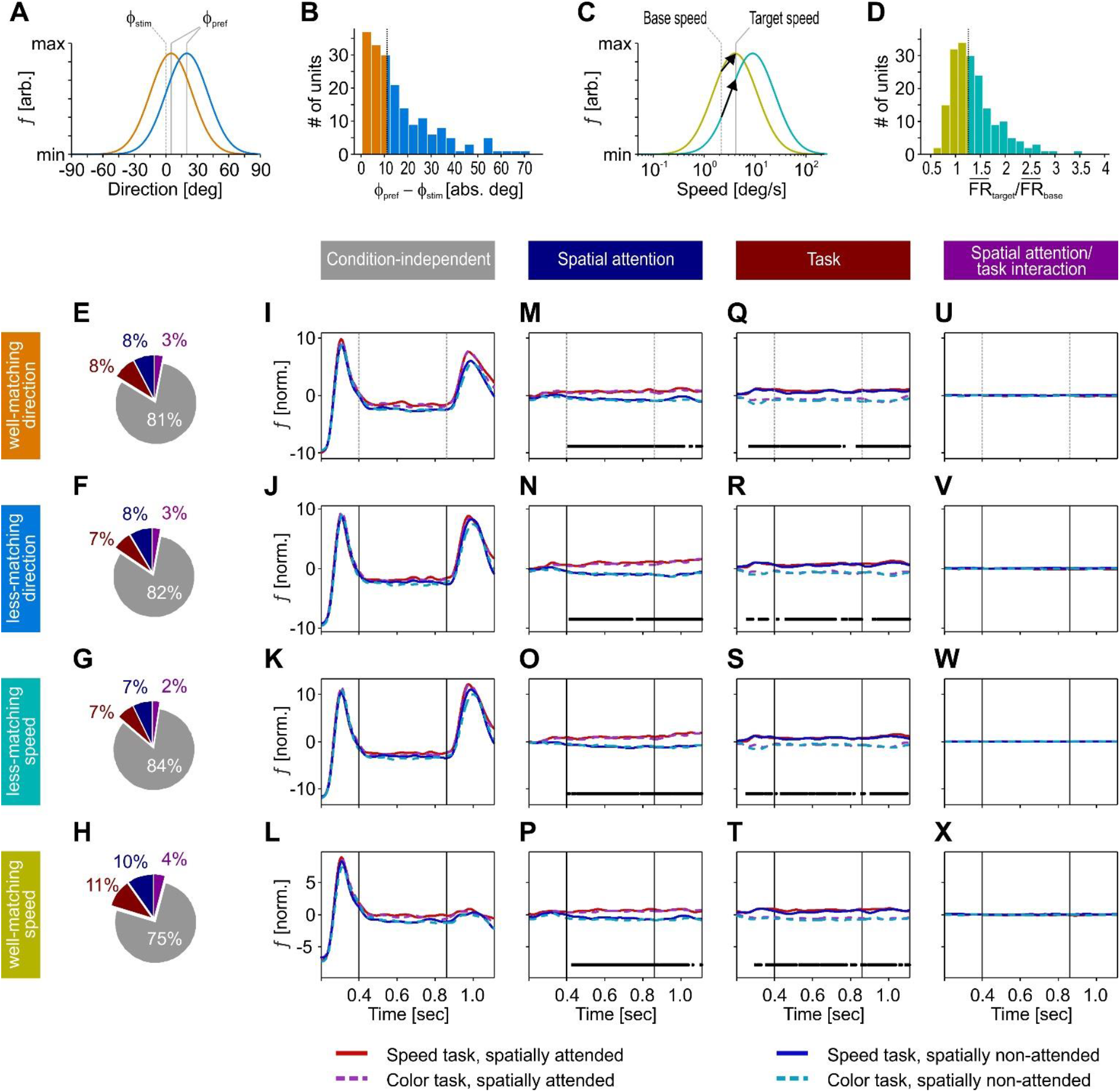
Influence of motion and speed preference on task-dependent response modulation. *A*: Sketch of two direction-tuning curves with small and larger deviation between preferred direction ϕ_pref_ and stimulus direction ϕ_stim_. *B*: Distribution of neurons depending on the absolute deviation ϕ_pref_ - ϕ_stim_, split by median (straight vertical line). Orange bars, well-matching neurons; blue bars, less-matching neurons. *C*: Sketch of two log-Gaussian shaped speed-tuning curves. Yellow line, speed-dependent responses of a neuron for which the base speed of the stimulus (dashed vertical line) is close to preferred and target speed (straight vertical line) is preferred; green line, speed-dependent responses of a neuron preferring higher speeds; black arrows, vectors indicating the FR increase for a jump from base to target speed, as expected from each neuron’s speed tuning. Note that the to-be-expected FR change of the sub-optimally driven neuron is about twice the size of the well-driven neuron. *D*: Distribution of transients with different amplitude, as expressed by the ratio between the post-change response and the pre-change response (test epochs 3 and 2 in Fig. 2), split by median (straight vertical line). Yellow bars, putatively well-matching neurons; blue bars, less-matching neurons. *E*-*H*: Percentage of explained variance for neuronal populations sorted by either direction preference or speed preference, estimated by dPCA. Colors in pie plots correspond to variance explained independent of condition (grey), by spatial attention (blue), task condition (red), and interaction between spatial attention and task (purple). *I* - *L*: Reconstructed spike trains showing explained variance essentially independent of condition, due to the passage of time. Colored lines correspond to the four experimental conditions, as indicated at the bottom of the figure. *M* - *P*: Reconstructed spike trains showing explained variance due to spatial attention, *Q* - *T*: task condition, and *U* - *X*: interaction of spatial attention and task. Black lines indicate periods of significant classification.

### Modulation of spontaneous activity

To investigate this dimension-based facilitation of MT responses in more detail, we compared the spontaneous activity before onset of the spatial cue. During this epoch, monkeys already knew the task type due to the color of the fixation point, but not the to-be-attended motion direction of the upcoming target. We found that FRs in the speed task were increased by 8.3% (Friedman test, *χ*^2^ = 43.46, *P* < 10^−10^, *N* = 187), while no such differences showed up if the same trials were sorted according to their later spatial condition (*χ*^2^ = 1.97, *P* = 0.16). We then investigated whether this baseline shift affected neurons differently, depending on their tuning (Fig. 5). We found, however, that neurons of all tuning groups were more active in the speed task. Note that regarding direction tuning, monkeys could not build an expectation about the upcoming target motion direction, and only knew that motion instead of color was relevant. The fact that both well-tuned and less-tuned neurons had a significant increase in baseline activity supports the hypothesis that in the speed task, MT received an early, general boost targeting neurons independent of their tuning. In contrast, monkeys could well anticipate the relevant speed, since this was the same inside and outside the spatial focus of attention. Yet, even neurons with the largest transients (for which the stimulus base speed was clearly away from their preferred speed) showed the same baseline increase than putatively well-tuned neurons. Neither in the speed nor in the color task, there was any statistical difference between tuning groups (Kruskal Wallis tests, both *χ*^2^(3) < 1.87, *P* > 0.59), but for each tuning group the across-task comparison revealed a highly significant baseline shift (Wilcoxon signed rank tests, all *Z* > 3.17, all *P* < 0.0015). This suggests that baseline activity in MT is actively adjusted as a function of task requirements.

**Fig. 5.**
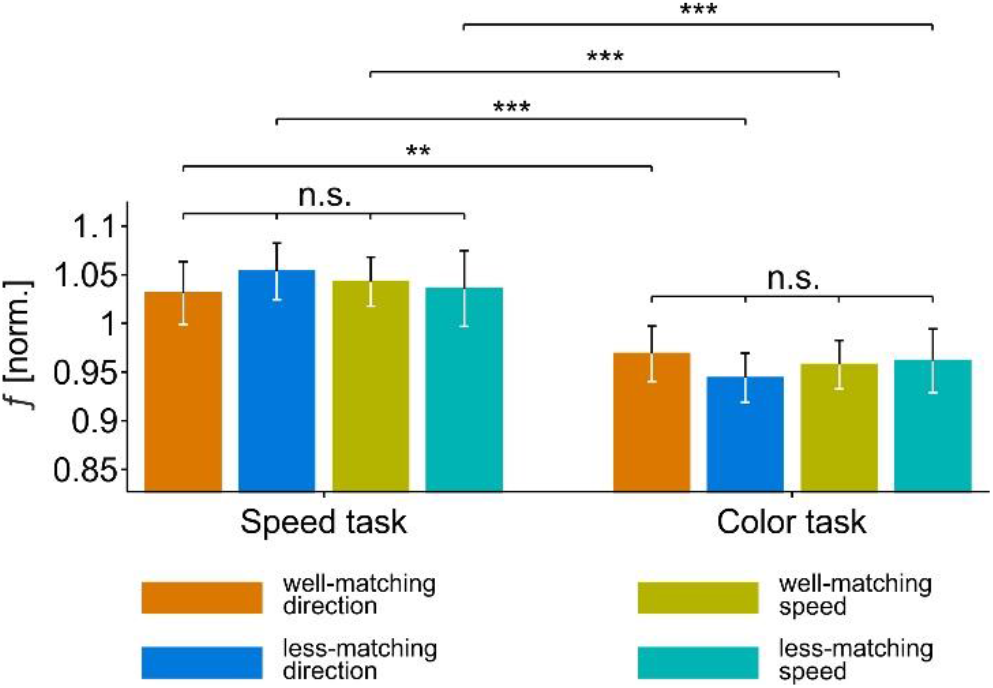
Normalized spontaneous activity depending on direction and speed-tuning preferences. Errors bars, 95% CI; *: *P* < 0.05, **: *P* < 0.01, ***: *P* < 0.001; n.s.: not significant.

To finally test whether this difference in baseline activity can account for the significant response differences during evoked responses, we subtracted the spontaneous activity in a condition-specific manner, i.e. we set all neurons to the same baseline level. Yet, this procedure did not eliminate the task-dependent modulation seen in the population SDF during visually evoked responses (Friedman test, *χ*^2^ = 12.7, *P* < 10^-3^, *N* = 187; *post-hoc* Wilcoxon signed rank tests inside and outside spatial focus of attention: *Z* > 2.97, *P* < 0.029), suggesting that task-dependent differences in the evoked response do not constitute a pure reflection of an early baseline shift but get significantly amplified during later visual processing.

## Discussion

Location-independent, feature-directed attention plays a crucial role in everyday life. A considerable amount of our knowledge about its neuronal mechanisms has been gained by experiments examining the processing of feature attributes, e.g. a specific color hue or motion direction (David et al. 2008; McAdams and Maunsell 2000; Treue and Martínez Trujillo 1999) and several models of visual attention built on attribute-specific top-down modulations (Corchs and Deco 2004; Wagatsuma et al. 2013). A key finding of studies on attribute-specific FBA is that attending to a specific motion direction facilitates the response of those neurons in area MT well-tuned to that attended attribute, while it does not modulate, or even suppresses, the response of neurons with a clearly different tuning (Martínez Trujillo and Treue 2004). These works constituted an important step forward in understanding the neuronal mechanisms by which the brain shapes information processing according to the behavioral requirements. Yet, there is another line of evidence suggesting a presumably more general modulation of neuronal responses when attending a specific stimulus feature. In a neuroimaging study requiring attention to either color or speed, both the baseline and stimulus-evoked BOLD signal were found to be enhanced in areas V4 and MT, respectively, depending on task requirements (Chawla et al. 1999). Because the signal increase was evident in MT even when subjects were presented with a stationary stimulus at the beginning of a speed-trial, and in V4 when they were presented with a monochromatic stimulus at the beginning of a color-trial, the authors concluded a task-dependent change of the attentional set, making those neurons more sensitive that process the relevant feature dimension. Similarly, in several visual search experiments, Müller and colleagues found that both reaction times and error rates benefit from a feature dimension-match between a preceding cue and the to-be-searched item. Interestingly, if the cue also matched the target’s feature attribute they did not find a further improvement of behavioral performance (Akyüek et al. 2010; Gramann et al. 2010; Töllner et al. 2010). These behavioral data were accompanied by cue dimension-related modulations of the event-related potential (ERP), which were evident both in the early component of the ERP indicating differences in attentional set, and in a later component reflecting a sustained, feature dimension-related attentional modulation. Finally, in a recent EEG experiment we were able to disentangle the influence of dimension- and attribute-specific attentional effects (Gledhill et al. 2015). Using a delayed match-to-sample task we found that a match in the feature dimension between a pre-cued, upcoming target stimulus and a spatially unattended, task-irrelevant stimulus is associated with a stronger negativity during the selection negativity-period (representing an event-related measure of feature selection (Anllo-Vento and Hillyard 1996; Torriente et al. 1999)), as compared to a stimulus defined in the non-attended dimension. A match in the feature-attribute provided an additional modulation on top of the dimension-based modulation, coming with longer latency and restricted to occipito-parietal electrodes.

The neurophysiological results described in the present study are well in line with these human psychophysical, EEG, and imaging studies, and suggest that task-specific, dimension-based effects of feature-directed attention are reflected by an attention-dependent weighting of the cortical module most sensitive to process the attended feature dimension. Besides an increase of response rates in the motion task (both with and without visual stimulation), MT neurons showed reduced trial-to-trial fluctuations and reduced spike-count correlations, and faster change transients when motion was behaviorally relevant, regardless of the spatial focus of attention. Our analyses showed that i) the task-specific FR differences during both baseline and evoked response are independent from the specific tuning of a neuron to the speed or direction of the motion stimulus, ii) the differences in the evoked response rate exceed the differences during baseline, and they still are of significant magnitude after full elimination of baseline-related FR differences, iii) the task-specific latency shortening of change transients is independent from both spatial attention and the relation between the attended and the stimulating motion direction, as to be expected from psychophysical results (Wegener et al. 2008; 2014), and iv) the differences in response variability and in response latency cannot be accounted for by differences in the firing rate, as they were evident also after rate-matching. Taken together, the results provide evidence for the notion of an attention-dependent, feature dimension-based weighting of visual processing at the neuronal level, and they provide mechanistic insights into earlier reports from human studies in support of an early dimension-based modulation of the attentional set and lasting modulatory processes influencing the processing of the behaviorally relevant stimulus (Chawla et al. 1999; Gramann et al. 2010).

### Comparison to other neurophysiological studies investigating task-specific components of visual attention

At the level of single cells, two recent studies addressing motion processing as a function of task requirements did not find corresponding experimental evidence ( Chen et al. 2012; Katzner et al. 2009). When recording from neurons in area MT, Katzner et al. (2009) investigated attentional modulations in a speed and color task, with an experimental paradigm similar to the one used in the present study. In accordance with our results, they found that spatial attention increases MT firing rates even if the anticipated stimulus change was a color change. Yet, the FSG-related response difference between attending a preferred vs. a non-preferred motion direction was of the same size irrespective of the task. The authors concluded that response modulation by both FBA and spatial attention occur independent of specific task requirements. Their results match the predictions of, and experimental findings from, object-based attention studies, proposing that attending a specific feature of a target object facilitates the processing of other, non-attended features of that object (Blaser et al. 2000; Duncan 1984; Ernst et al. 2013; O'Craven et al. 1999; Rodríguez et al. 2002; Schoenfeld et al. 2003; Wannig et al. 2007). Another MT study required monkeys to make a saccade into one of four previously trained directions in response to either the color or the motion direction of a briefly presented grating stimulus (Chen et al. 2012). In 25% of the trials, the saccade direction for indicating the color were congruent with the motion direction of the grating, but in 75% of the trials they were incongruent, requiring the animal to focus on the task at hand to provide the correct response. Yet, only 22% of MT neurons showed significantly different responses between tasks, and of these, some were more active during the direction task, and others were more active during the color task.

Despite methodological differences, including experimental paradigms, visual stimulation, and data analysis, we consider one major difference in the experimental design to be important to understand the proper conditions under which the visual system facilitates or suppresses the processing of an entire feature dimension. In both former studies, monkeys had to frequently switch between motion and color trials, while in the present study they were required to engage in one type of task for several dozen trials. A strong engagement in one specific task type (and hence, strong attention to one specific feature dimension) is, however, more likely to occur if this task is performed for a prolonged time, as compared to frequently switching between tasks. This interpretation is in line with psychophysical investigations using similar task and stimulus conditions as in the present neurophysiological experiments (Wegener et al. 2008). In this study, subjects were required to respond to any speed or color change as fast as possible. When they were given a feature dimension cue of 75% validity, they responded very quickly to the indicated change, but much slower if the change occurred in the unattended feature dimension, regardless of the spatial focus of attention. In contrast, if they were given an object cue without specific information on the feature dimension most likely to change, reaction times were in-between the previous distributions, i.e. they were slower as compared to the previously attended feature dimension, but faster as compared to the previously unattended dimension. These data indicate that strongly attending to one feature dimension may go along with the suppression of processing in another dimension, preventing the spreading of attention to other, irrelevant features of the selected object, and thereby increasing behavioral performance. Similar results on the suppression of non-attended object features have been obtained in several other studies (Cant et al. 2008; Fanini et al. 2006; Freeman et al. 2014; Nobre et al. 2006; Polk et al. 2008; Serences et al. 2009; Taya et al. 2009; Xu 2010). As such, a possible reason for the different results on the task-dependency of MT responses is that in our experiments monkeys were more strongly focusing on the feature relevant for the current task block. In contrast, monkeys in the previously described studies may have been allocating their attention more to the entire object than to a specific feature, resulting in object-based spreading of attention to task-irrelevant features and less suppression of the irrelevant feature-dimension in Katzner et al. (2009), and to a significantly reduced performance for incongruent stimuli in Chen et al. (2012).

### Results cannot be explained by feature-similarity gain

Our finding of a tuning-independent, feature dimension-specific response modulation is in conflict with the FSG hypothesis, which constitutes the current and most influential mechanistic explanation of FBA. This conflict does not arise from a general incompatibility with FSG (since FSG may co-exist with the task-specific modulation we here describe), but from the fact that FSG cannot account for the response differences as found in our experiments. Specifically, attending to the non-preferred motion direction outside the RF is not expected to enhance but rather suppress the FR of the recorded neuron (Martínez Trujillo and Treue 2004). Because of the dissimilarity between the attended motion direction outside the RF and the stimulating, preferred motion direction inside the RF, we concluded that the response difference between the color and the speed task cannot be explained by an attention mechanism based on the similarity between attended and preferred motion direction.

A possible concern, however, to our interpretation of a tuning-independent modulation may arise from the following assumption: If it was possible to solely attend the speed but not the direction of the target Gabor, FSG potentially facilitates the response of neurons tuned to the attended speed, which might be stronger if speed changes have to be detected instead of color changes. If a significant portion of our neurons would prefer the speed of the stimulus before the change (which was the case), then these neurons may be addressed by FSG, and the observed task-specific response differences could be interpreted within the FSG framework without assuming a tuning-independence of feature-directed attention.

We argue, however, that this interpretation is unlikely and does not explain the discrepancy between previous findings on FSG and the results of the current study. First, this alternative interpretation still depends on a task-specific response difference. As explained earlier, the effect of FSG was found to not vary between tasks, such that FSG is interpreted to address those feature-specific neurons that process the attended object (Katzner et al. 2009). Thus, even if most of our neurons preferred the speed prior to the change, based on the findings of Katzner et al. (2009) FSG would have the same effect in both tasks. This conclusion is not supported by our results. Second, conceptually the selective facilitation of only those neurons well-tuned to the speed before the change is unlikely to support change detection. In a speed-change task, neurons tuned to the attended speed are the poorest change detectors within the population of speed-tuned neurons (Traschütz et al. 2015). Because they respond to their preferred speed, any change in this speed, be it an acceleration or a deceleration, results in a reduction of their response. Due to the log-Gaussian shaped speed tuning curves of most MT neurons (Lagae et al. 1993; Nover et al. 2005), these neurons produce very small transients even for noticeable changes, which tend to average out in the population (Traschütz et al. 2015). In contrast, neurons for which the attended speed is on the rising or falling part of their tuning curve respond with significant FR changes even to small changes in speed, and preserve the sign of the speed change (acceleration or deceleration) over a large spectrum of speeds. Experimentally, we recently showed that the response of a population of neurons with heterogeneous speed tuning profiles (such as the population response we measured in the current study) is mainly carried by such neurons, having a preferred speed away from the stimulating speed (Traschütz et al. 2015, Fig. 4). Hence, suppressing the response of these neurons while facilitating the response of neurons with a preference to the attended speed hardly serves to improve change detection. In accordance with this and discouraging an interpretation based on speed tuning-profiles, we found that third, dimension-based attentional modulation is addressing neurons independent of their speed tuning (Fig. 4), and even in the absence of visual stimulation (Fig. 5). As such, we consider a selective modulation of only those neurons tuned to the target speed as not capable to explain the experimental findings of the current study.

### Mechanistic implications for feature-directed attention

In our previous EEG experiment (Gledhill et al. 2015), we found that feature dimension-specific attentional effects emerged at frontal electrodes and then moved over to parieto-occipital electrode sites over visual cortex. In the present study, the first noticeable difference between the speed- and the color task was a global shift in the baseline activity of MT neurons, prior to stimulus onset, which suggests a different attentional set (Corbetta and Shulman 2002) in the motion task. In line with the EEG findings (Gledhill et al. 2015), a possible source for such task-related top-down modulations of visual cortical activity is the prefrontal cortex (PFC) (Miller and Cohen 2001). Neurons in monkey PFC have a different degree of motion- and color sensitivity depending on the behavioral relevance of either feature, and color- and motion-selective neurons cluster in different parts of the PFC (Lauwereyns et al. 2001. Lesions to the lateral PFC impair shifting of the attentional set to another perceptual dimension (Dias et al. 1996). Therefore, our result of a task-dependent increase in baseline activity is in line with the notion of an attention-related biasing signal from attentional control areas to specialized visual modules (Chawla et al. 1999; Driver and Frith 2000).

Although baseline shifts are not sufficient on its own to induce attention-related response differences during the evoked response (Fannon et al. 2008), and do not predict the magnitude of attentional modulation (McMains et al. 2007), they often precede both spatial and feature-based attentional modulations in visual cortex (Chawla et al. 1999; Chelazzi et al. 1998;Kastner et al. 1999; Lee et al. 2007; Luck et al. 1997; Reynolds et al. 2000; Shulman et al. 1999). In line with this, we found that both the feature dimension and spatial attention effects were not dependent on the baseline increase observed during the pre-stimulus epoch, but had a significant magnitude after subtraction of spontaneous activity, even if done for each of the four behavioral conditions separately.

Taken together, the current data show that directing attention between motion and color of a stimulus causes (at least under the task- and stimulus conditions of our experiment) a task-specific, dimension-based attentional modulation of neuronal activity. Unlike FSG, this modulation is not only addressing the neurons that are well-tuned to the attended stimulus but all neurons in the cortical module most sensitive to process the attended stimulus dimension.

## Acknowledgements

The work was funded by DFG Grants KR 1844/1-2 and WE 5469/2-1, and two ZF Grants from the University of Bremen. The authors declare no competing financial interests. The authors acknowledge support by Peter Bujotzek, Sunita Mandon, Deniz Parmuk, Ramazani Hakazimani, and Katrin Thofe regarding various aspects of the study.

Current address of MP: School of Medical Sciences, University of New South Wales, Australia.

## Author contributions

Conceptualization: D.W. and A.K.K.; Methodology: D.W., A.K.K., B.S., and F.O.G.; Software: F.O.G., B.S., A.K.K., and D.W.; Formal Analysis: B.S. and D.W.; Investigation: B.S., F.O.G., and M.P.; Writing - Original Draft: D.W. and B.S.; Writing - Review & Editing: D.W. B.S., and A.K.K.; Funding: D.W. and A.K.K.; Supervision: D.W. and A.K.K.

